# A frozen human wound-repair reference reveals heterogeneous tumor niches and conditional immunotherapy associations

**DOI:** 10.64898/2026.09.10.750782

**Authors:** Qi Wu, Shuiming Jiang, Lei Liu

## Abstract

**Purpose:** Tumors are often framed as unresolved wounds, but the extent to which independently defined physiological repair programs are recontextualized across cancers remains uncertain.

**Methods:** We derived five lineage-associated modules and a composite score from 58,823 cells in a longitudinal human skin-wound cohort before examining tumors. Frozen modules were evaluated in paired hepatocellular carcinoma and lung adenocarcinoma single-cell datasets, spatial and cell-resolved non-small-cell lung cancer cohorts, and four pretreatment immune-checkpoint-inhibitor cohorts. Patient- or section-level inference and prespecified sensitivity analyses were retained.

**Results:** Myeloid repair modules showed the most consistent tumor-associated increase in paired cancers; other lineage effects varied by context. Repair-high non-small-cell lung cancer regions combined epithelial/proliferative, extracellular-matrix fibroblast, and C1QC/broad-myeloid programs, whereas repair-cytotoxic neighborhood correlations were split across sections. In six Prime 5K Xenium patients, repair-high regions were fibroblast enriched (+0.239 high-minus-low fraction; 6/6 concordant; exact sign-flip P=0.031; FDR=0.047) and T/NK depleted (−0.151; 6/6; FDR=0.047). Direct ligand-receptor neighbor effects were not reproducible. Repair_FBI showed an adverse ICI-response direction, but the random-effects estimate was imprecise (OR=1.69, 95% CI 0.81-3.51).

**Conclusion:** Tumors reuse elements of physiological repair to form heterogeneous multicellular niches. Fibroblast remodeling is the most stable candidate; the evidence supports conditional association, not universal immune exclusion, a senescence-defined state, or a generalizable ICI biomarker.

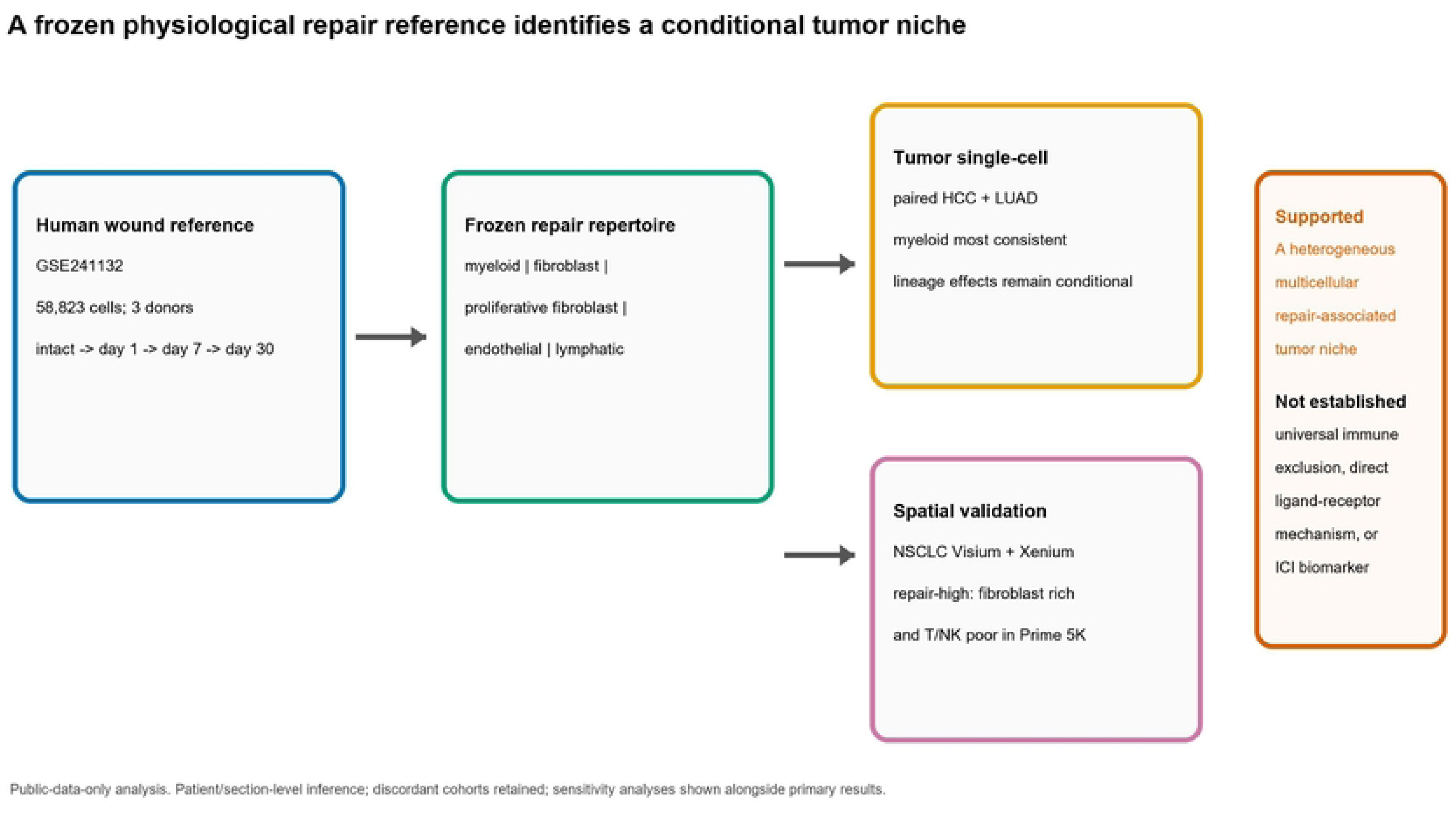

## 1. Introduction

The analogy between cancer and a wound that does not heal has informed studies of stromal activation, extracellular-matrix remodeling, angiogenesis, and immune recruitment for decades [1,2]. Physiological repair, however, is not a single transcriptional state. It is a time-ordered process in which epithelial, inflammatory, fibroblast, endothelial, and lymphatic programs overlap and resolve differently across tissues [3,4]. Repair and cancer can share lineage plasticity without being identical processes [19]. Tumor microenvironments are likewise composed of heterogeneous malignant, stromal, vascular, and immune cell states [5–8]. A broad “wound-like” score may consequently represent different cell mixtures, spatial arrangements, or technical sampling properties in different cancers.

Single-cell and spatial profiling have made this distinction experimentally addressable. They reveal heterogeneous cancer-associated fibroblast and myeloid states and demonstrate that regional tumor-immune architectures can have divergent clinical implications [5,6,9–12]. At the same time, a lineage-localized score, a repair-high regional composition, a ligand-receptor prediction, and an ICI association are distinct evidentiary claims. None by itself establishes causality or a predictive biomarker. Pan-cancer microenvironment classifications have also identified recurring states while underscoring disease-context dependence [13].

We therefore used a staged design in which a physiological human wound cohort served exclusively for module construction. Scores and gene weights were frozen before analysis of tumors, spatial topology, or ICI outcome. Independent data were assigned non-overlapping roles: paired tumor-normal datasets tested tumor recontextualization; spatial datasets tested regional architecture; cell-resolved spatial data tested composition and short-range neighbor hypotheses; and treatment cohorts tested translational association. The objective was to identify the reproducible core and context-dependent limits of a wound-repair repertoire, without converting association into a causal “tumor hijacking” or myeloid-senescence claim.

## 2. Materials and methods

### 2.1. Study design and public datasets

This public-data, multi-stage study used discovery, validation, spatial-architecture, clinical-association, and robustness cohorts separately (Table 1). The final interpretation emphasized effect direction, patient/section concordance, score coverage, and replication role rather than the smallest P value. No additional stringent cell-level filter was introduced beyond the data structures provided by the original submitters. Individual cells were not treated as independent patients for tumor-normal, regional-composition, or treatment-response inference. Expression matrices and metadata were obtained from Gene Expression Omnibus (GEO) [14].

**Table 1.** Public datasets and prespecified roles.

| Evidence stage | Dataset | Disease/context | Unit of inference | Prespecified role |
| --- | --- | --- | --- | --- |
| Physiological discovery | GSE241132 | Human skin wound healing; 58,823 cells, 3 donors | Patient-time-cell-type pseudobulk | Module construction and freezing only |
| Tumor validation | GSE149614 | Paired HCC tumor/non-tumor | Patient-cell-type pseudobulk | Cross-cancer tumor recontextualization |
| Tumor validation | GSE131907 | Paired lung adenocarcinoma | Patient-cell-type pseudobulk | Independent paired direction check |
| Spatial architecture | GSE238264; GSE292299 | Treated HCC; treatment-naive NSCLC Visium | Section | Spatial support and regional topology |
| Cell-resolved spatial | CosMx Lung9; GSE311609 | NSCLC CosMx/Xenium | Patient after section aggregation | Neighbor boundary test and composition |
| Clinical association | GSE78220; GSE91061; GSE126044; GSE135222 | ICI-treated melanoma/NSCLC | Patient | Context-dependent translational association |

Dataset roles were fixed before results were synthesized. The wound cohort was not reused for cancer validation, and an ICI cohort was not used to tune a response score. Patient was the replicative unit in paired single-cell analyses; sections were retained rather than selectively pooled in Visium analyses; and Xenium sections were first summarized within patient. This hierarchy limits precision inflation from many cells or repeated sections from the same donor and permits discordant components to remain visible.

### 2.2. Physiological discovery and module freezing

GSE241132 included 58,823 submitter-filtered human skin-wound singlet cells from 12 samples, three donors, and four time points: intact skin and days 1, 7, and 30 after injury. Temporal changes were summarized by patient-by-time-by-cell-type pseudobulks. We retained complete descriptive rankings rather than imposing a hard discovery threshold. Five 150-gene positive wound-associated modules were created: Repair_MonoMac, Repair_FBI (fibroblast injury/remodeling), Repair_FBProlif_D7, Repair_VE_D7, and Repair_LE. A 300-gene Repair_Composite was derived by union ranking. Module membership and gene weights were frozen before examination of tumor or ICI labels.

External scores were calculated on observed genes and renormalized for module coverage with a rank-based signature-scoring framework [15]; unmeasured genes were not set to zero. The composite is intentionally broad and was used to describe a mixed regional niche. Component-level analyses were always retained because broad scores can be sensitive to shared proliferation or low panel coverage.

Module construction retained positive wound-associated rankings rather than selecting only genes passing a stringent adjusted-P-value threshold. The goal was to encode a physiological repertoire and then assess transferability, not to maximize separation in a three-donor discovery dataset. Downstream gene-removal, unweighted, coverage-aware, and component-deletion analyses address the consequences of this inclusive choice. Sensitivities were parallel evidence and were never used to remove an unfavorable primary cohort.

### 2.3. Paired tumor single-cell analyses

GSE149614 was analyzed as paired HCC tumor/non-tumor data and GSE131907 as paired primary lung adenocarcinoma tumor/non-tumor data. Patient-cell-type pseudobulks were created for lineages with at least 15 cells in the relevant specimen. Exact paired sign-flip testing was the primary directional test. Limma-voom gene-set testing, expression-matched random-gene-set controls, and alternative cell-count thresholds were complementary robustness analyses. Concordance was evaluated by lineage and module, rather than with a pooled pan-cancer P value.

### 2.4. Spatial analyses and cell-resolved composition

GSE238264 treated HCC Visium data and GSE292299 treatment-naive NSCLC Visium data were scored with frozen repair and fixed cytotoxic programs. The primary topology statistic was the within-section Spearman correlation between a spot’s repair score and the mean cytotoxic score of its six nearest neighbors. Four- and eight-neighbor graphs were prespecified sensitivities. In GSE292299, repair-high and repair-low regions were compared using epithelial/proliferative, extracellular-matrix fibroblast, C1QC/broad-myeloid, vascular, and lymphocyte state programs. These scores were interpreted as local transcriptional states, not deconvolved cell fractions. Cell-resolved analyses used CosMx Lung9 and GSE311609 Xenium. CosMx neighbor testing preserved field-of-view structure and applied within-field, marker-state-conditioned permutations. In GSE311609, all coordinate-matched positive-library cells from 22 NSCLC sections were retained. Donor-balanced centroid transfer from a public author-labeled lung reference supplied Level-3 labels; marker-dominant broad labels were retained as an annotation sensitivity, consistent with the limitations of reference-dependent integration [16]. Prime 5K sections from six patients were the primary composition analysis because they measured 119 of 300 composite genes. Custom immune-oncology sections measured 23 composite genes and were analyzed only as lower-coverage sensitivity data. Repair-high and repair-low cells were the upper and lower score quartiles within each section. Section estimates were aggregated to patients and assessed with exact sign-flip tests; FDR was calculated across prespecified cell-label tests.

The distinction between Prime 5K and lower-coverage panels was prespecified at synthesis. A targeted panel that lacks much of the 300-gene composite cannot be assumed to measure the same state as a high-coverage panel. We therefore retained the lower-coverage data as platform and annotation sensitivities rather than pooling them as equivalent validation. For every spatial modality, a positive repair-neighborhood association was interpreted as a local association, not as proof of intercellular communication.

### 2.5. Clinical association and sensitivity analyses

Four pretreatment ICI cohorts were retained: GSE78220, GSE91061, GSE126044, and GSE135222. Deposited clinical response definitions were used; progression-free survival was assessed where available. No score cutpoint was trained. Cohort-specific response estimates were displayed together, and a random-effects summary was descriptive because disease, assay, endpoint, and sample-processing heterogeneity were expected. The terminal sensitivity analysis was parallel to, not a replacement for, the primary analysis. It included GSE126044 fresh-tissue restriction, removal of proliferation-associated or candidate ambient-RNA genes, unweighted scoring, leave-one-component-out composite scoring, and alternative response definitions when metadata allowed. Spatial proximity was not interpreted as signaling: direct ligand-receptor hypotheses required reproducible state-conditioned local-neighbor effects, given known autocorrelation and expression-state confounding in spatial communication analyses [17,18].

## 3. Results

### 3.1. A frozen wound-repair repertoire captures temporal multicellular remodeling

The human wound time course showed staged organization rather than a monotonic high-repair state (Fig. 1). Early samples showed epithelial expansion. Day 7 was characterized by monocyte/macrophage, fibroblast injury/remodeling, proliferative-fibroblast, and vascular endothelial programs, while day 30 was relatively fibroblast dominant. This temporal and lineage-resolved structure motivated the five component modules and the composite. Because the reference was frozen before cancer analyses, subsequent tumor and ICI results could not redefine module membership or gene weights.

**Fig. 1.**
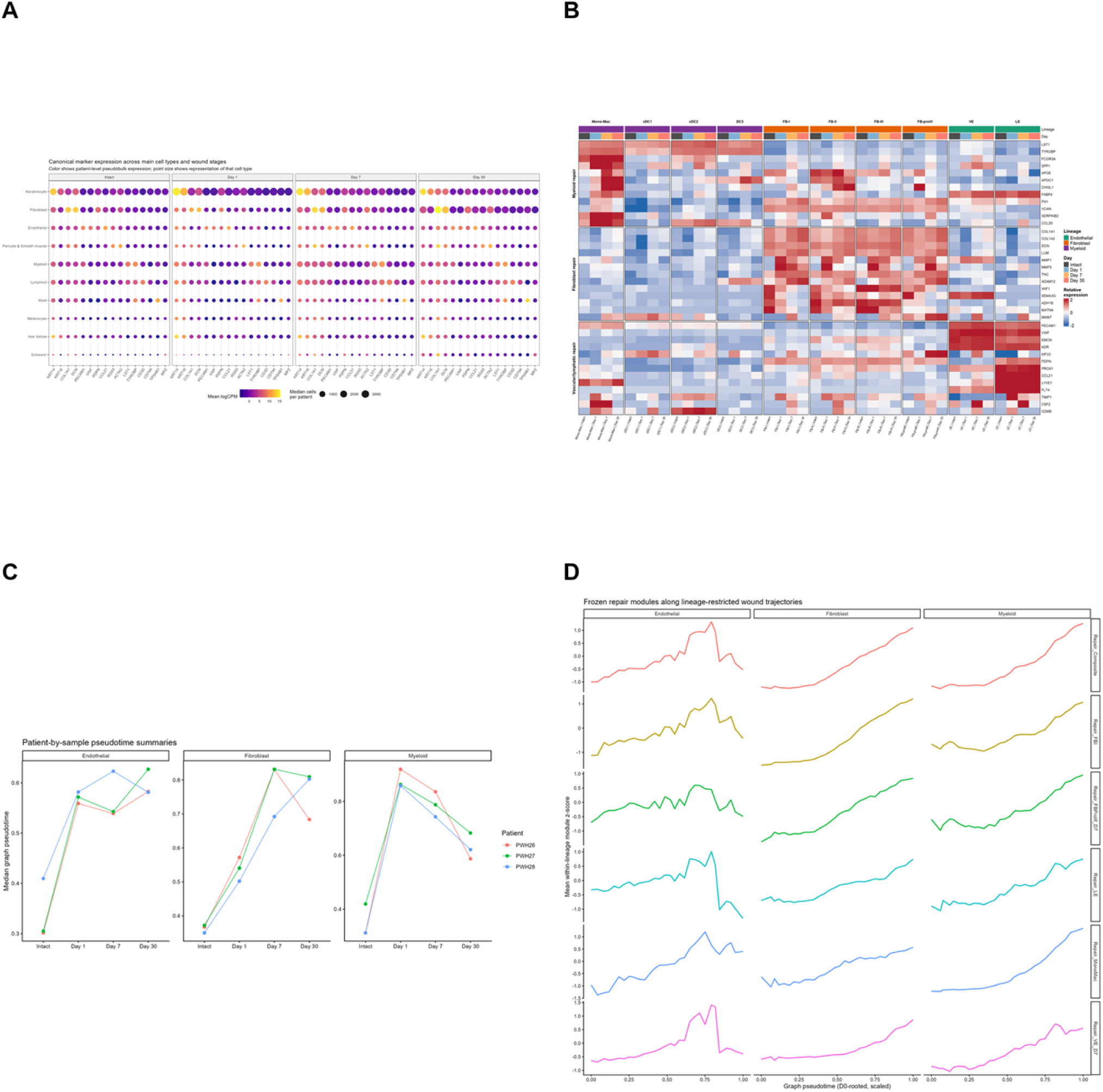
Human physiological wound healing defines a frozen multicellular repair repertoire. (A) Cell-type marker and composition overview across intact, day-1, day-7, and day-30 wounds. (B) Subtype/time-course expression heat map. (C) Patient/sample-resolved temporal organization. (D) Module curves along temporal ordering. The repertoire was frozen before tumor and ICI analyses.

### 3.2. Paired tumors show a reproducible myeloid/fibroblast core with context-dependent extensions

In paired HCC, the six frozen modules localized to expected but non-exclusive lineages (Fig. 2A). All six myeloid module directions were positive in patient-paired tumor-normal pseudobulks (6/6 concordant directional observations; exact sign-flip P=0.008). Fibroblast injury/remodeling was also coherent across complementary statistical approaches, whereas proliferative-fibroblast and endothelial components were more method dependent (Fig. 2B). The conclusion is not that all tumor myeloid cells share one state; it is that the wound-associated myeloid arm is the most reproducible tumor-associated continuation of this physiological reference in HCC.

**Fig. 2.**
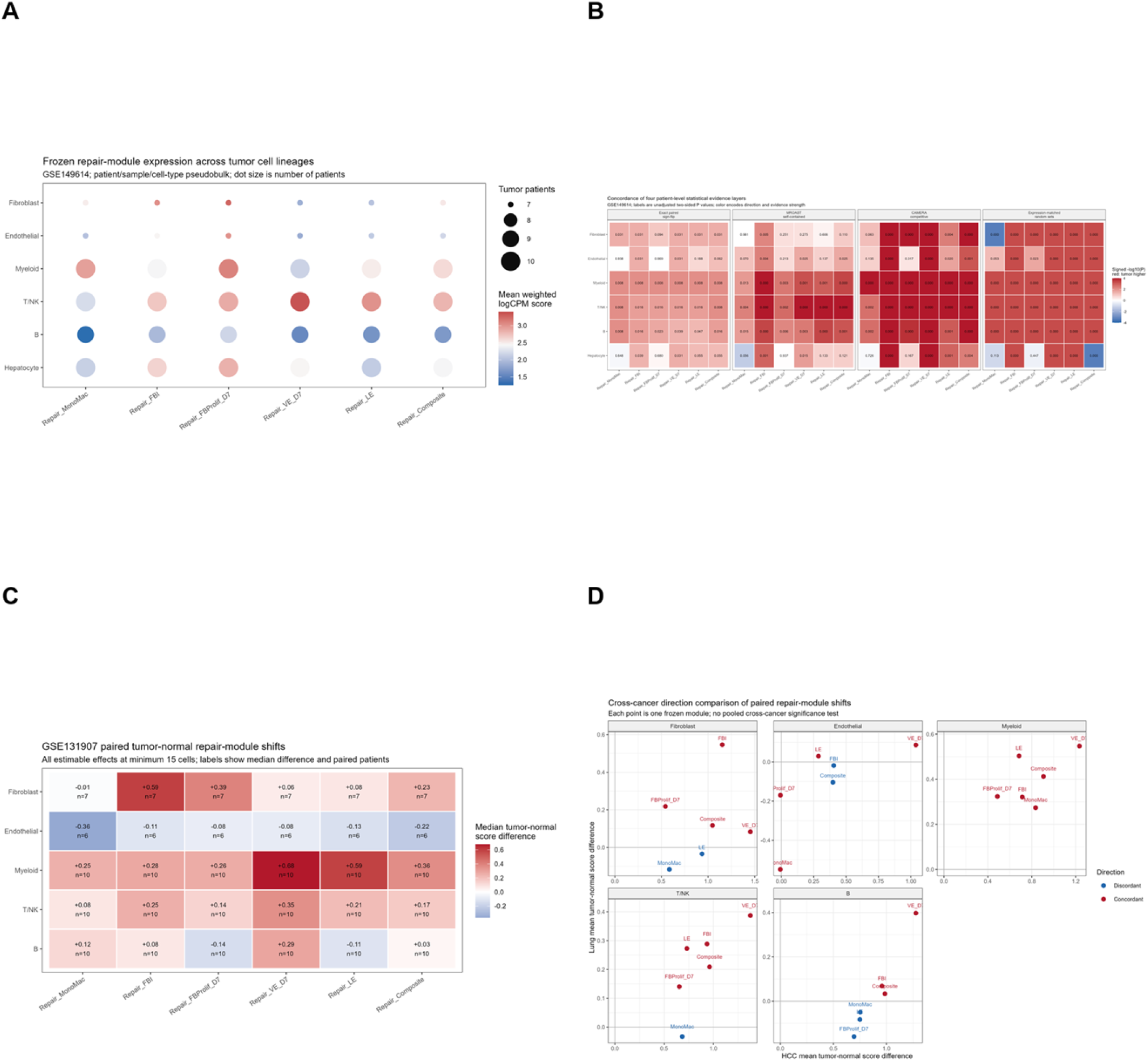
Paired tumor datasets identify a recurrent myeloid/fibroblast repair core. (A) HCC lineage localization of six frozen modules. (B) Patient-paired HCC tumor-normal effects across complementary approaches. (C) Independent paired lung adenocarcinoma effects. (D) HCC-lung direction concordance. Patient-level pseudobulks are the inferential unit.

The independent lung cohort supported a restricted cross-cancer core. Twenty-two of 30 lineage-by-module directions agreed between HCC and lung, including six of six myeloid component directions. Repair_FBI was positive in seven paired lung comparisons. Endothelial and monocyte/macrophage-associated patterns were not uniformly aligned across cancers (Fig. 2C-D). Thus, the paired analysis supports a recurrent myeloid/fibroblast repair axis, but does not support pan-lineage or pan-cancer elevation of every component.

This agreement was sufficient to reject a purely sparse-cell interpretation, but not homogeneous enough to justify one pan-cancer effect size. Expression-matched random-gene-set controls and alternate pseudobulk cell-count thresholds preserved the dominant HCC myeloid/fibroblast pattern without converting weaker endothelial or proliferative components into positive findings. The lung dataset consequently acts as a directional external check rather than as an artificial replication of the HCC lineage mixture.

### 3.3. Repair-high regions are mixed niches with conditional cytotoxic topology

Treated HCC spatial data provided supporting evidence of repair/cytotoxic colocalization in selected contexts. Treatment-naive NSCLC was therefore the principal regional-architecture cohort. Repair-high regions combined epithelial/proliferative, extracellular-matrix fibroblast, and C1QC/broad-myeloid states (Fig. 3). This pattern extends the paired single-cell results by showing that the repertoire is a regional multicellular niche rather than a single-lineage signal.

**Fig. 3.**
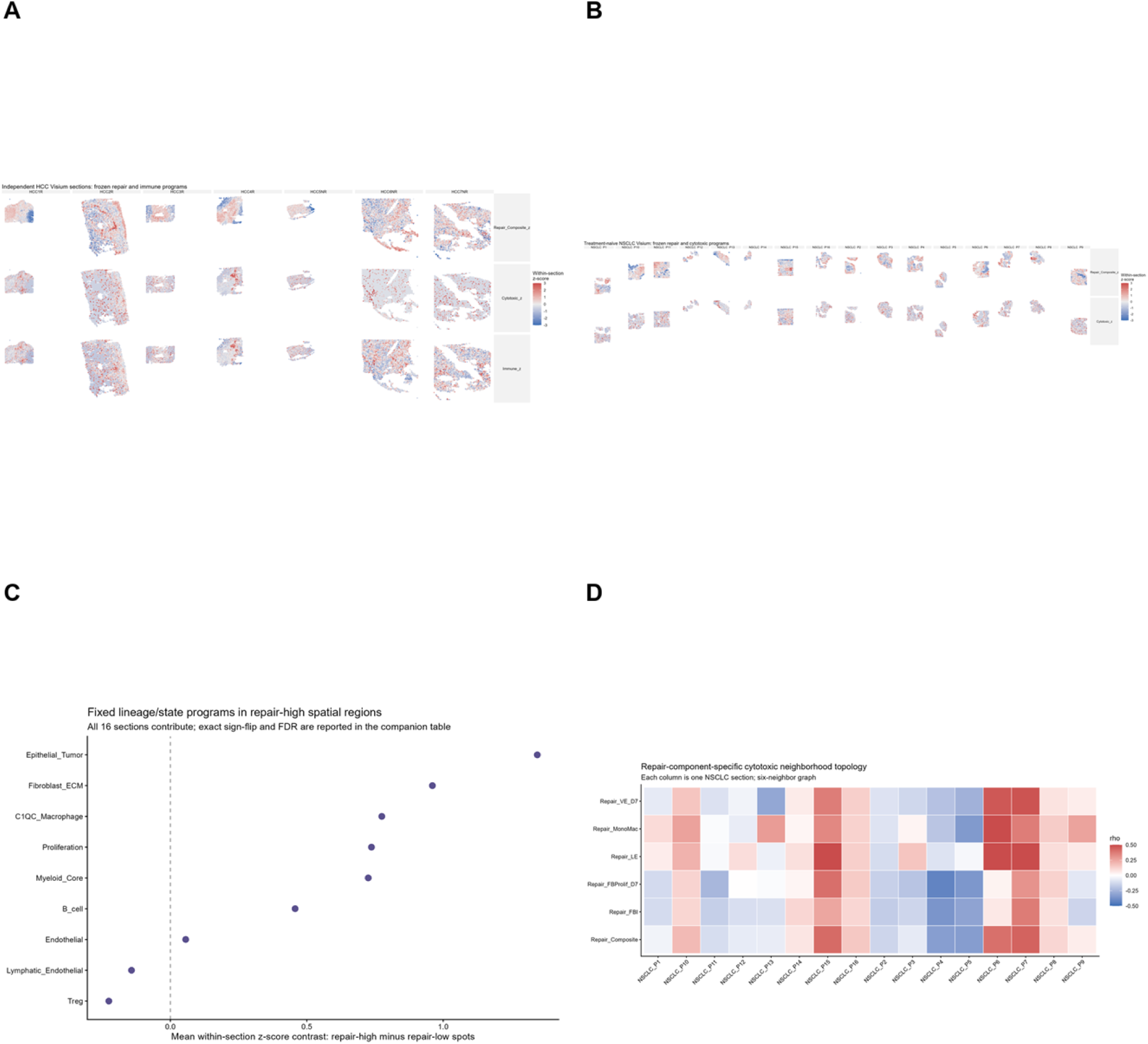
Repair-high spatial regions form a mixed niche with conditional cytotoxic topology. (A) Treated HCC repair/cytotoxic maps. (B) Treatment-naive NSCLC repair/cytotoxic maps. (C) Repair-high versus repair-low regional state contrasts. (D) Component-specific repair-to-cytotoxic neighborhood associations. State scores are not deconvolved cell fractions.

Repair-high regions did not show a universal cytotoxic-exclusion topology. In treatment-naive NSCLC, repair-to-cytotoxic neighborhood correlations were evenly split across sections (eight positive and eight negative) and remained qualitatively mixed with four-, six-, and eight-neighbor graphs. Component-level behavior also varied. Consequently, the appropriate interpretation is conditional topology, not universal exclusion of cytotoxic cells. Treatment context further constrained interpretation. Treated HCC maps showed that repair and cytotoxic signals can colocalize, but they could not determine a treatment-naive general rule. The treatment-naive NSCLC cohort provided that test and showed that a fibroblast/myeloid-rich region does not invariably form a cytotoxic desert. Retaining this mixed result prevents regional composition from being incorrectly recast as a universal immune-exclusion mechanism.

### 3.4. High-coverage Xenium data support repair-associated fibroblast enrichment and T/NK depletion

In GSE311609, reference transfer and the original marker-dominant annotation agreed for several broad categories but were not interchangeable (Fig. 4A). Both were retained to prevent post hoc selection of the annotation that favored a desired effect. The Prime 5K panel supplied the most interpretable composite coverage and formed the primary composition analysis.

**Fig. 4.**
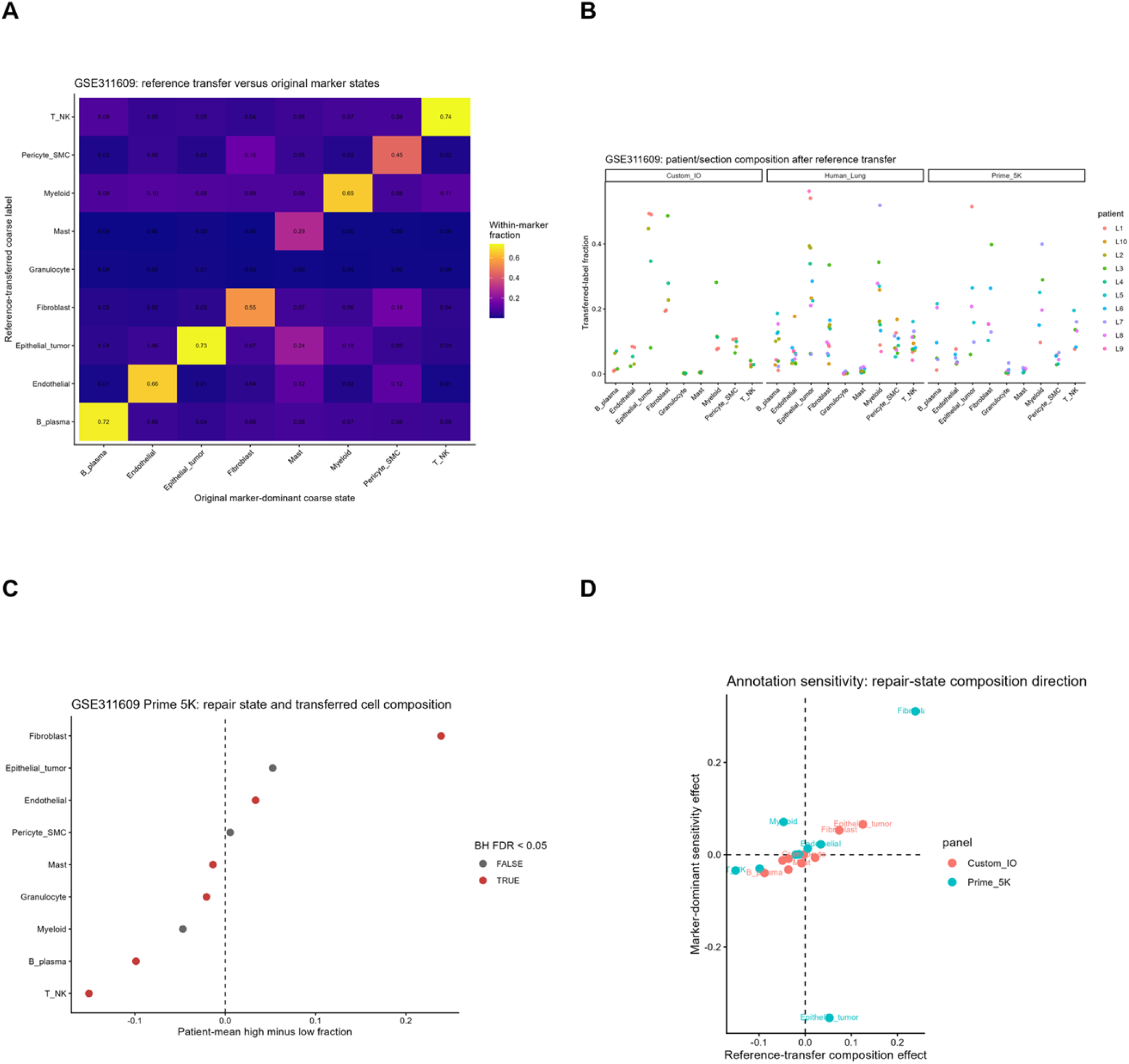
Cell-resolved spatial data support repair-associated fibroblast enrichment and T/NK depletion. (A) Reference-transfer versus marker-dominant annotation agreement. (B) Composition after reference transfer. (C) Primary Prime 5K patient-level high-minus-low composition effects. (D) Annotation sensitivity. Fibroblast enrichment and T/NK depletion were each concordant in six of six patients; direct ligand-receptor neighbor effects were not reproducible.

Across six Prime 5K patients, repair-high regions had higher fibroblast fractions than repair-low regions (+0.239 patient-mean high-minus-low fraction; six of six patients positive; exact sign-flip P=0.031; FDR=0.047) and lower T/NK fractions (−0.151; six of six patients negative; FDR=0.047). B/plasma fractions were also lower (−0.099; FDR=0.047); endothelial effects were small (+0.034; FDR=0.047), and epithelial direction was unstable. Fibroblast enrichment persisted under marker-dominant annotation (+0.311; six of six patients), although the sensitivity-family FDR was 0.070 (Fig. 4B-D). Direct state-conditioned ligand-receptor neighbor effects were not reproducible. Therefore, regional composition is supported in the primary high-coverage panel, whereas a specific direct signaling mechanism is not.

The patient-level concordance is critical because the raw data contain many cells and multiple sections. The fibroblast and T/NK effects were assessed from six donor-level high-minus-low contrasts, rather than from a cell-level P value. Conversely, six patients do not establish a universal NSCLC property. We therefore present this as the most specific spatial composition result in the project, with a defined panel-coverage and patient-number boundary.

### 3.5. ICI associations are component-specific and imprecise

All ICI cohorts were retained in the final display (Fig. 5). GSE78220 showed supportive adverse directions for Repair_FBI (area under the receiver operating characteristic curve 0.798; odds ratio 3.37) and the composite (odds ratio 3.57), with borderline multiple-testing adjustment. GSE91061 was weak, null, or opposite under the deposited response definition; alternative disease-control definitions altered the apparent direction. In GSE126044, fresh-only analysis (n=11) retained adverse Repair_FBI and vascular-endothelial directions despite fresh/formalin-fixed paraffin-embedded composition. GSE135222 progression-free survival effects were weak and imprecise. Repair_FBI was the only component with an adverse direction in all three response cohorts. Its descriptive random-effects odds ratio was 1.69 (95% confidence interval 0.81-3.51), which is insufficient for a clinical biomarker claim. Composite and lymphatic effects attenuated after proliferation-gene removal, while myeloid effects were heterogeneous. Repair_FBI retained its adverse direction after candidate ambient-RNA-gene removal and unweighted scoring. Sensitivities therefore refine the evidence: fibroblast injury/remodeling is a prospective candidate, whereas composite and myeloid clinical interpretations remain conditional.

**Fig. 5.**
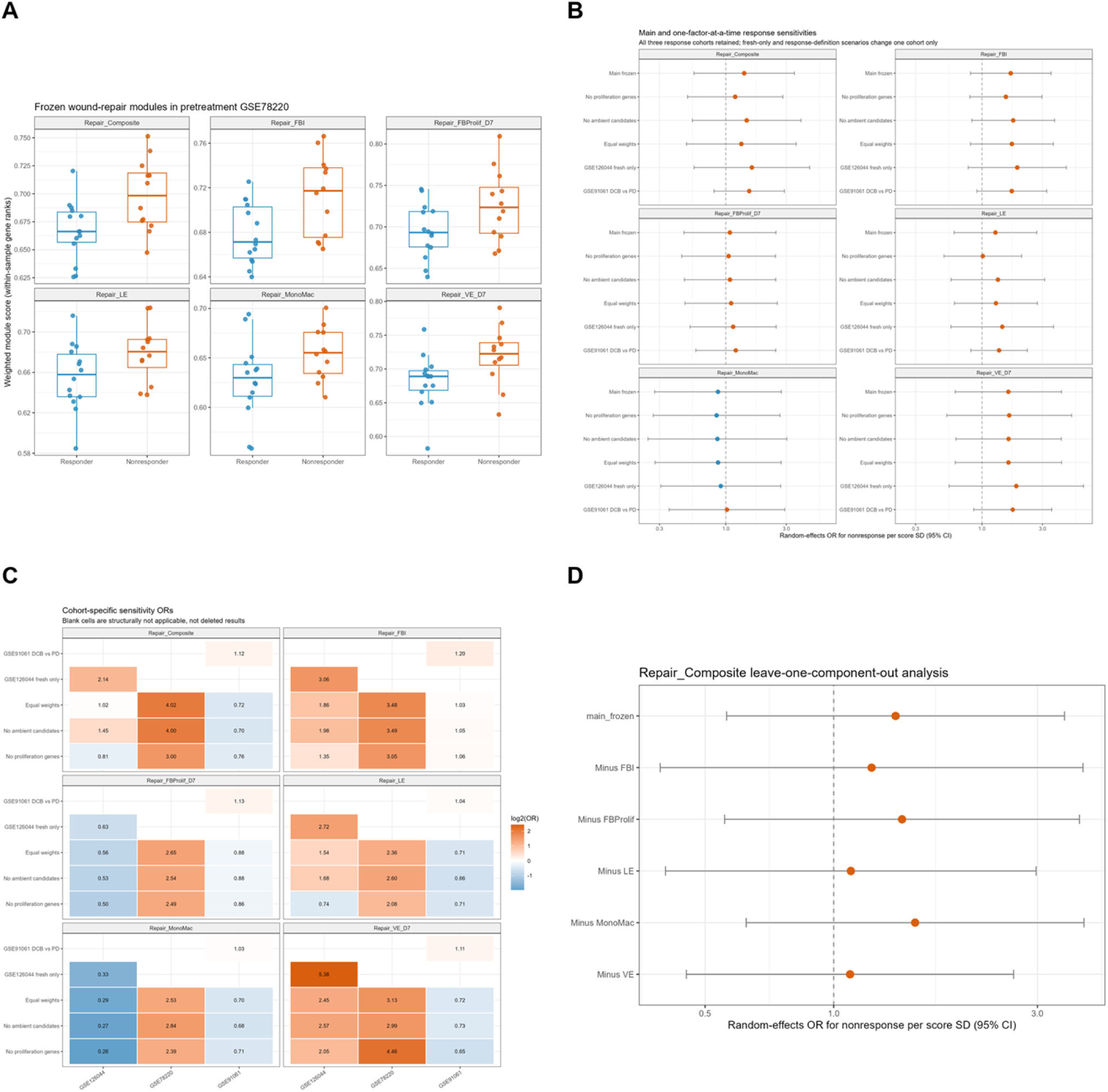
ICI associations are component-specific and interpreted with terminal sensitivity analyses. (A) GSE78220 response-associated score distributions. (B) Cohort-specific response effects and descriptive random-effects summary. (C) Cohort/sensitivity heat map. (D) Composite leave-one-component-out analysis. Fresh-only, gene-removal, unweighted, alternative-response, and component-deletion analyses are shown beside the primary results.

The sensitivity design identifies where the inference depends on operational choices. Fresh-tissue restriction addresses processing heterogeneity in GSE126044; gene-removal analyses identify components vulnerable to proliferation or ambient-RNA structure; and alternate response definitions expose endpoint dependence in GSE91061. Showing each result beside the primary estimate avoids selectively retaining only the most favorable clinical grouping or score specification.

## 4. Discussion

We used a physiological wound reference, frozen before tumor or outcome analysis, to identify a recurrent but heterogeneous repair-associated tumor niche. The strongest supported conclusion is a multicellular myeloid/fibroblast core. Its extension to endothelial, lymphatic, proliferative, and cytotoxic programs varies by lineage, cancer type, spatial scale, and score construction. In the primary high-coverage NSCLC Xenium analysis, repair-high regions were fibroblast rich and T/NK poor. In ICI cohorts, Repair_FBI showed a consistently adverse direction but remained too imprecise and context dependent for biomarker use.

The frozen design is important because it keeps biological definition separate from clinical outcome. It also makes component heterogeneity visible. Classical wound-like tumor descriptions often combine matrix, myeloid, vascular, and immune processes into a single stromal state [1,2,20]. Here, those components did not behave interchangeably: myeloid signals were most consistent in paired tumors, fibroblast injury/remodeling was the clearest regional and clinical candidate, and broad composite/lymphatic scores were sensitive to proliferation-gene removal. The composite is therefore useful for naming a mixed niche, but component-level reporting is necessary for interpretation.

Our spatial data distinguish regional composition from immune topology and direct signaling. The Prime 5K result supports a repair-associated composition shift, but the Visium repair-cytotoxic correlations were mixed and direct neighbor tests were negative. These observations can coexist because different technologies and spatial scales measure different features. They also argue against equating coexpression or adjacency with ligand-receptor communication. The absence of a reproducible direct local effect is an informative boundary result, not a negative experiment to hide.

The ICI analyses require similarly calibrated language. Repair_FBI is compatible with literature implicating fibroblast- and transforming-growth-factor-beta-associated states in immune restraint [21,22], but GSE91061 discordance and a pooled confidence interval spanning the null prevent a general response-biomarker claim. This study is complementary to, rather than a replication of, broader microenvironment classifications [13]. It nominates a fibroblast-remodeling component for prospective testing with homogeneous treatment cohorts and prespecified score coverage.

Negative mechanistic results further define the manuscript’s scope. Wound-time ordering provided moderate structure for fibroblast programs but did not reveal a universally ordered myeloid trajectory. Prior-based transcription-factor candidates lacked cross-context mRNA support, and prespecified spatial axes were not accompanied by reproducible direct-neighbor effects. These data do not rule out regulation or signaling in individual tumors; they do make a universal master regulator, virtual knockout phenotype, or accelerated physiological myeloid-senescence claim inappropriate here.

This study has limitations. The discovery cohort contains three donors; the modules contain shared injury and proliferation genes; Visium scores are regional transcriptional states rather than cell fractions; and the primary Xenium result relies on 119/300 composite genes in six patients. Reference transfer is model dependent, and lower-coverage panels should not be pooled as equivalent validation. All findings are observational. Perturbation, chromatin, spatial protein, and longitudinal treatment data will be required to test causal regulation or treatment vulnerability.

In summary, tumors reuse elements of physiological repair to form heterogeneous regional ecosystems. The current evidence supports a context-dependent repair-associated niche and prioritizes fibroblast injury/remodeling for prospective validation. It does not establish accelerated physiological myeloid senescence, a universal direct ligand-receptor mechanism, immune exclusion, or a generalizable ICI biomarker.

## Statements and Declarations

### Ethics approval and consent to participate

Ethics approval and informed consent were not required for this secondary analysis of publicly available, de-identified datasets. No participant contact, intervention, recruitment, or access to identifiable private information was involved. The original studies were conducted under the ethics approvals and consent procedures applicable to their respective source cohorts.

### Consent for publication

Not applicable. No individual-level identifiable information is reported in this article.

### Data availability

The primary datasets analyzed in this study are publicly available in the Gene Expression Omnibus under accession numbers GSE241132, GSE149614, GSE131907, GSE78220, GSE91061, GSE126044, GSE135222, GSE238264, GSE292299, and GSE311609.

---

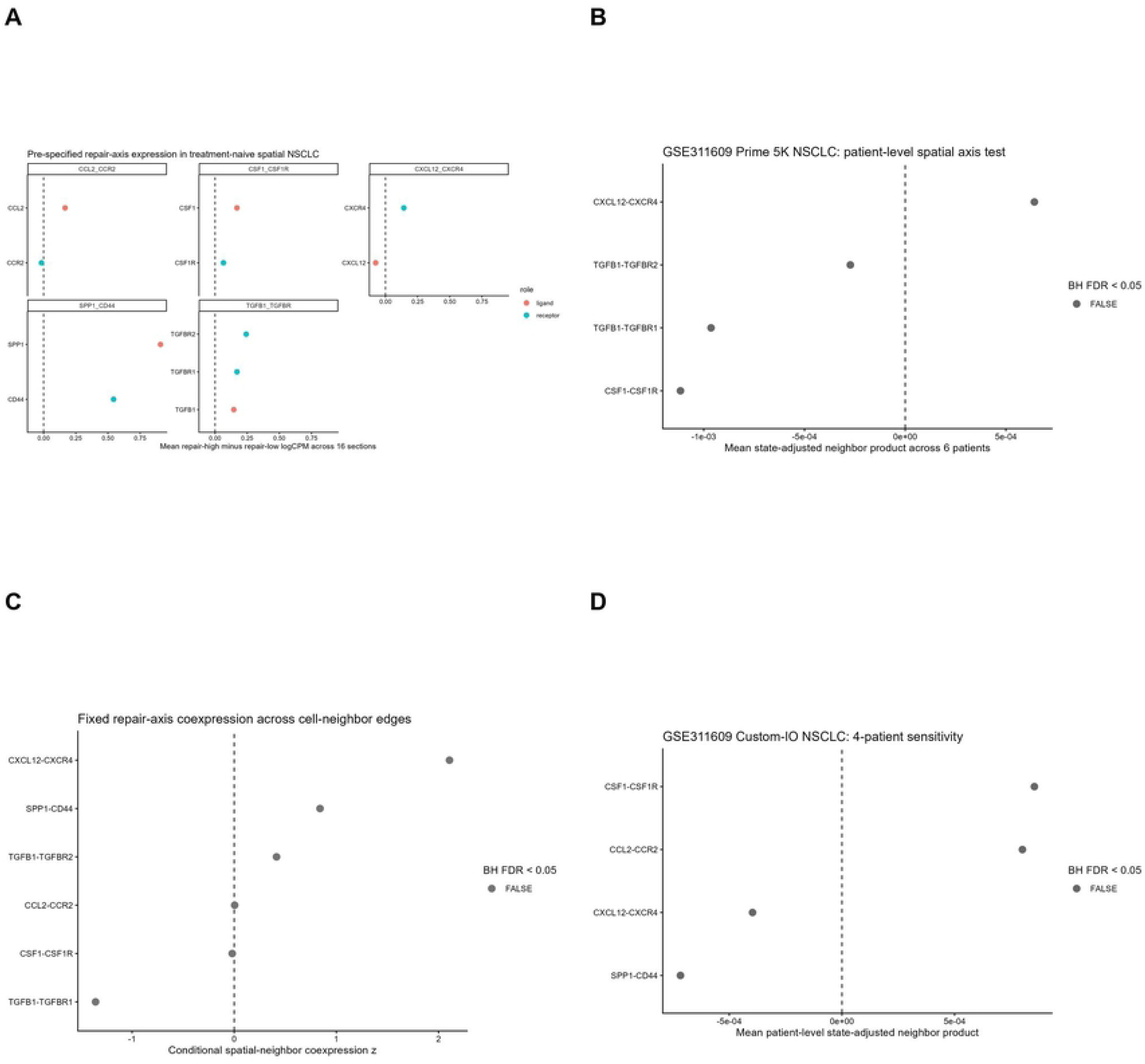

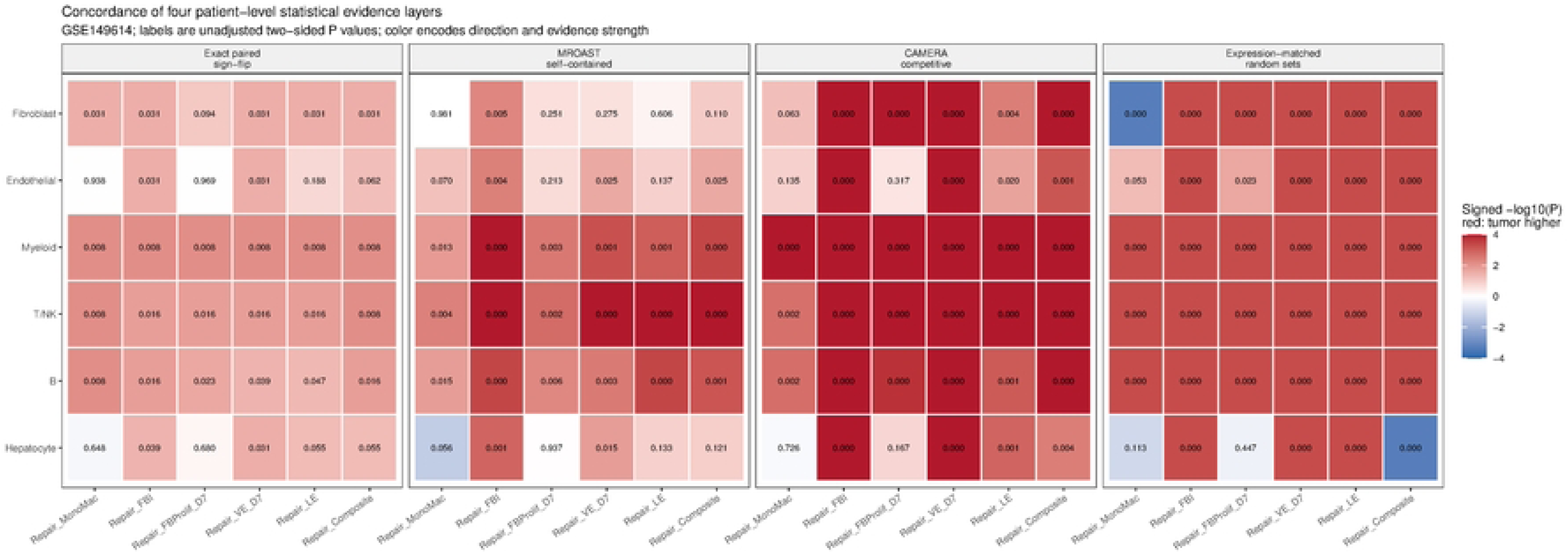

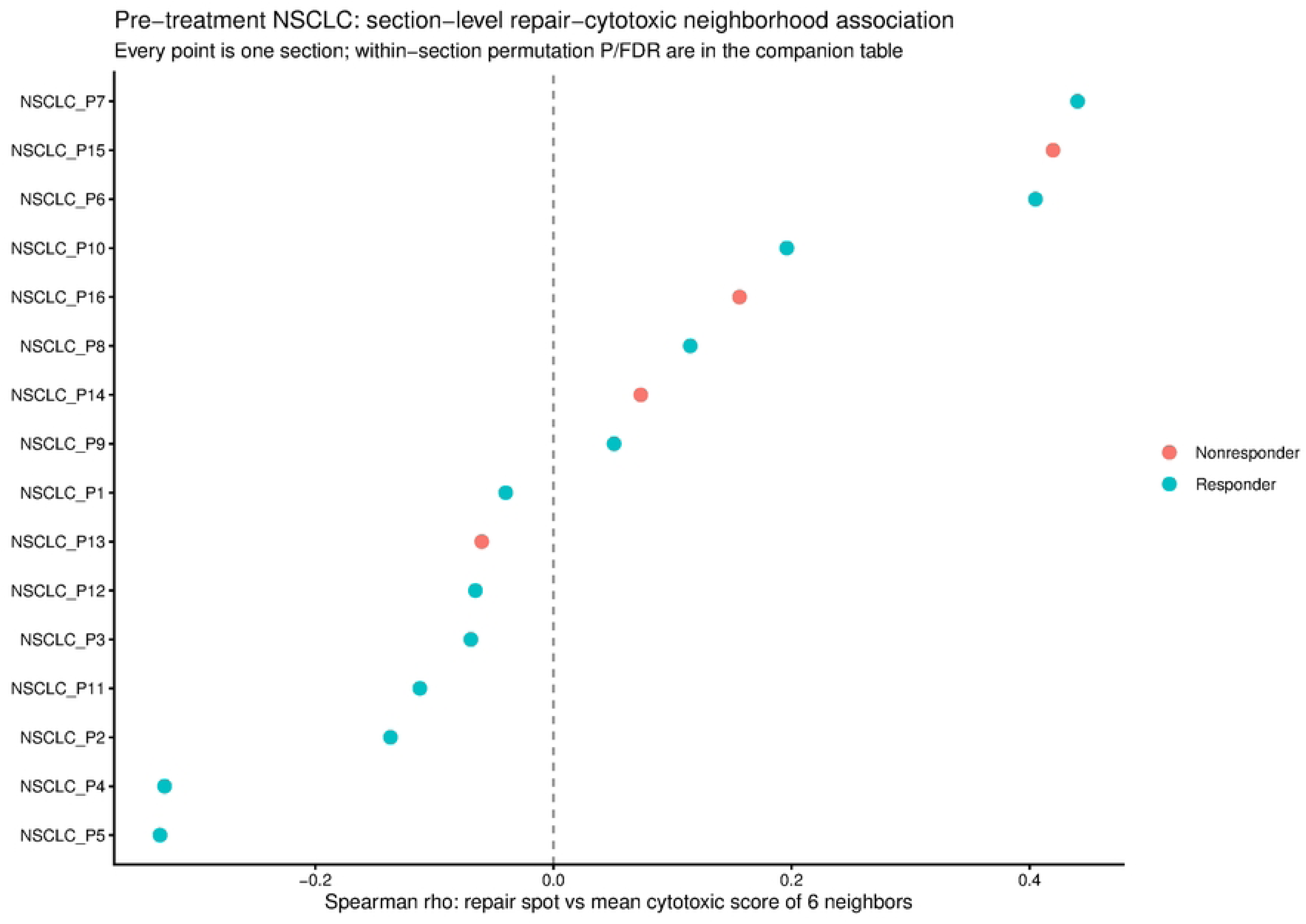

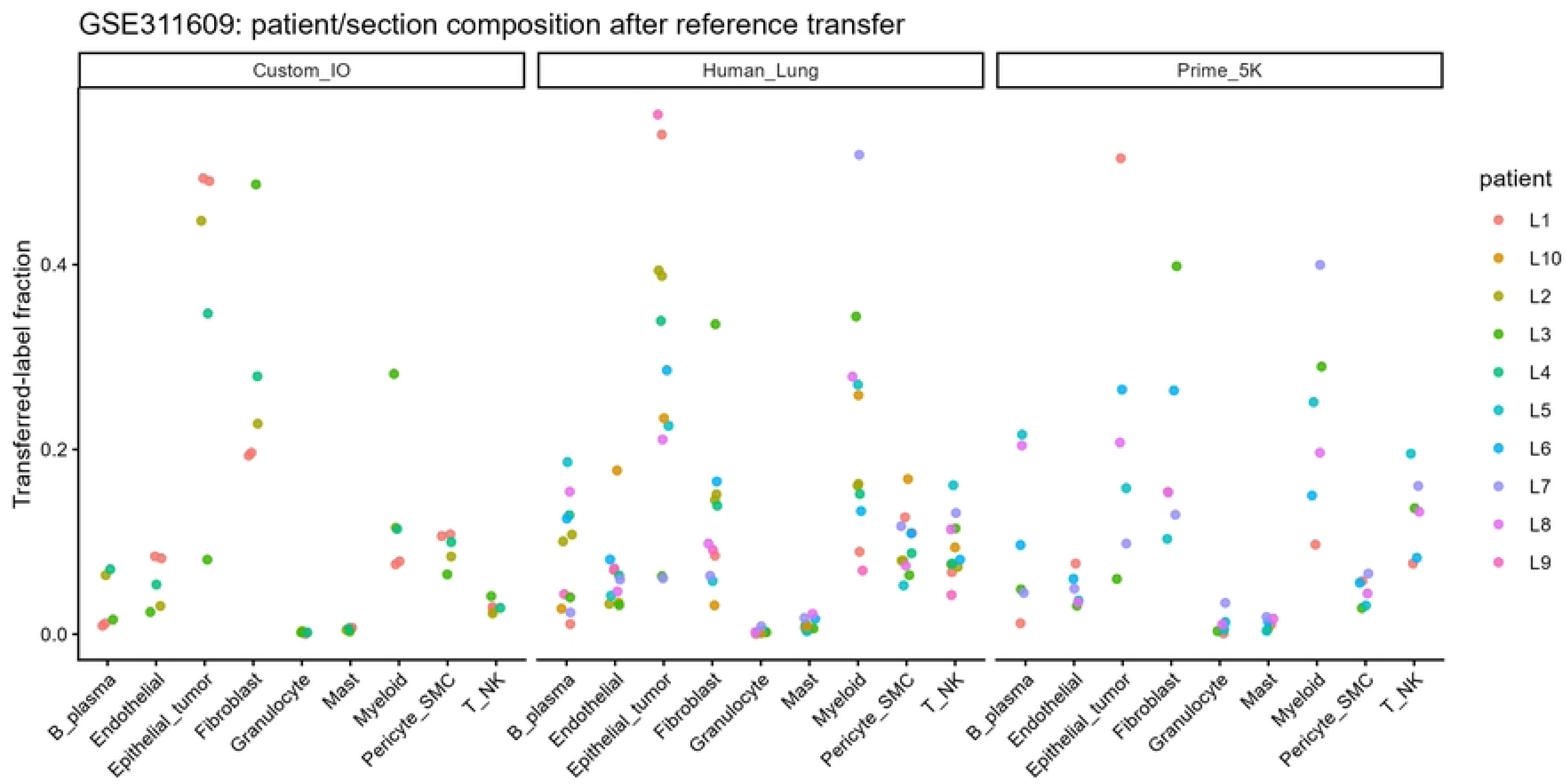

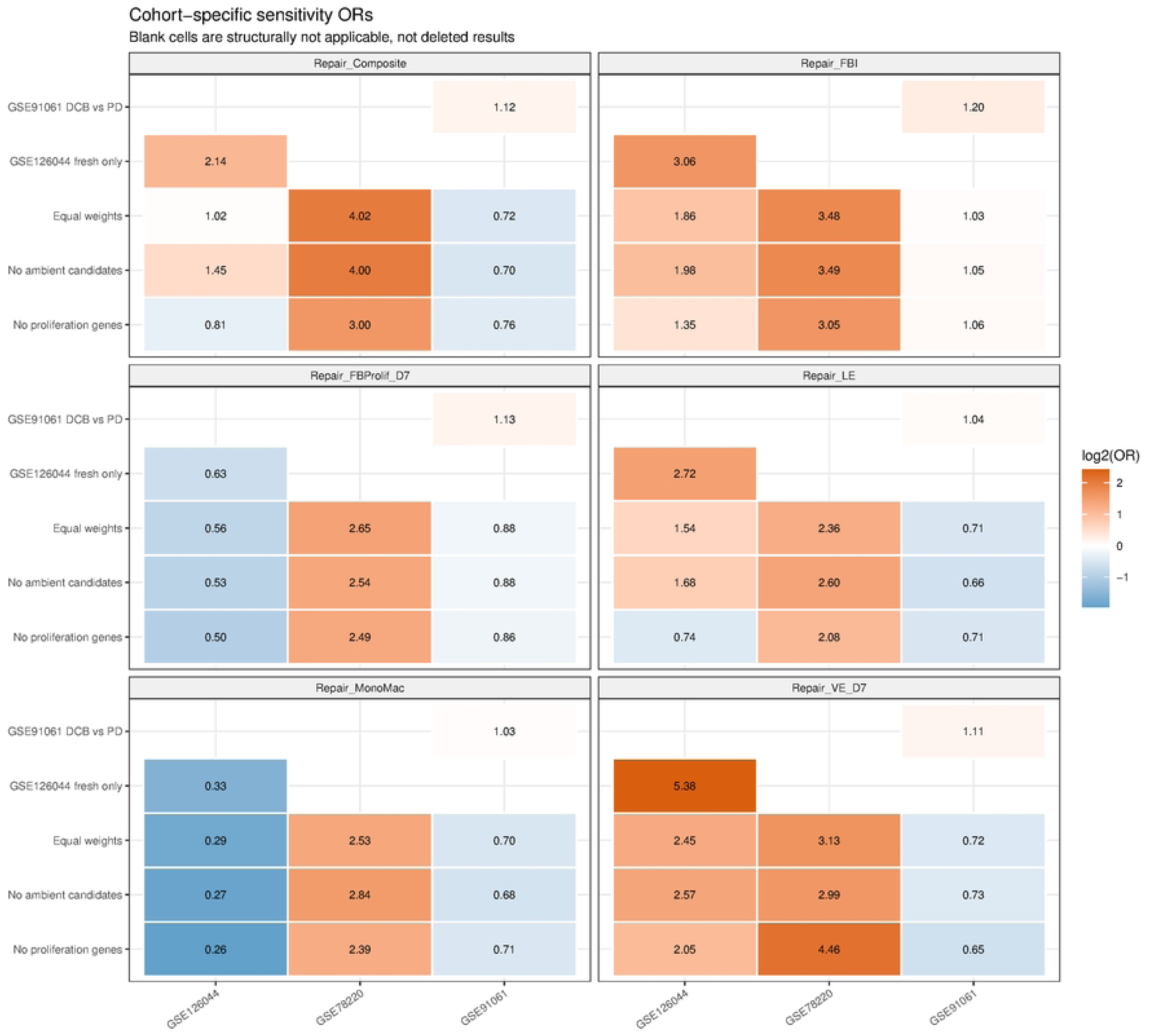

